# Variability in training unlocks generalization in visual perceptual learning through invariant representations

**DOI:** 10.1101/2022.08.26.505408

**Authors:** Giorgio L. Manenti, Aslan Satary Dizaji, Caspar M. Schwiedrzik

## Abstract

Stimulus and location specificity are long considered hallmarks of visual perceptual learning. This renders visual perceptual learning distinct from other forms of learning, where generalization can be more easily attained, and unsuitable for practical applications, where generalization is key. Based on hypotheses derived from the structure of the visual system, we test here whether stimulus variability can unlock generalization in perceptual learning. We train subjects in orientation discrimination, while we vary the amount of variability in a task-irrelevant feature, spatial frequency. We find that independently of task difficulty, this manipulation enables generalization of learning to new stimuli and locations, while not negatively affecting the overall amount of learning on the task. We then use deep neural networks to investigate how variability unlocks generalization. We find that networks develop invariance to the task-irrelevant feature when trained with variable inputs. The degree of learned invariance strongly predicts generalization. A reliance on invariant representations can explain variability-induced generalization in visual perceptual learning, suggests new targets for understanding the neural basis of perceptual learning in high-order visual cortex, and presents an easy to implement modification of common training paradigms that may benefit practical applications.

## Introduction

A fundamental problem for perception is to extract reliable information from a highly variable signal^1^. It is widely accepted that the visual system achieves this by learning what is consistent in its inputs, a process called *perceptual learning* (PL). However, how the enormous variability in the environment^2^ impacts PL *itself* is not well understood. Here, we ask how the visual system solves the challenge of variability for learning.

Variability poses both a problem and an opportunity for PL. Varying stimuli are problematic because they entail reduced predictability^3^, weaker memory traces^4^, uncertain reward assignment^5^, and non-linear decision rules^6^. They can thus slow down or impair learning. Consequently, state-of-the-art research and commonly employed PL protocols aim to maximize PL using highly unnatural conditions where variability is minimized, e.g., when learning one stimulus alternative at a time. However, what is often overlooked is that variability can be a great asset for learning as it facilitates *generalization* (i.e., applying learned behavior to new stimuli): variability may foster the extraction of core features across stimuli through abstraction, concept learning, and rule derivation. In many learning domains, variability acts as a catalyst for generalization^7^. In fact, theoretical studies suggest that the degree of variability during learning determines whether the system specializes on specific stimuli by memorizing them or instead learns generalizable rules^8^. Although generalization is frequently considered the ultimate goal of learning^9^, the current mainstream view on PL disregards variability and generalization. Instead, PL is often said not to generalize^10,11^. This renders visual PL an outlier amongst many learning phenomena.

Yet, the structure of the visual system suggests (at least) two ways to solve the challenge of variability during PL: the system could *generalize* by relying on invariant representations or *specialize* on both task-relevant and -irrelevant aspects of the stimuli, using highly precise neurons narrowly tuned to both dimensions. If the visual system employs any of these strategies is currently unknown.

A generalization strategy based on invariant representations, e.g., in ventral temporal cortex (VTC), deals with variability by subsuming it: invariant representations provide information about visual stimuli in an abstract form, irrespective of task-irrelevant variability in low-level details of the input (Fig. 1A). Thus, even if stimuli differ in appearance from trial to trial, invariant neurons still systematically provide relevant information, and can serve as a substrate for PL despite variability. However, these neurons are often not very precise: e.g., they signal which orientation is presented, but cannot distinguish orientations as accurately as neurons in early visual areas^12^.

**Figure 1:**
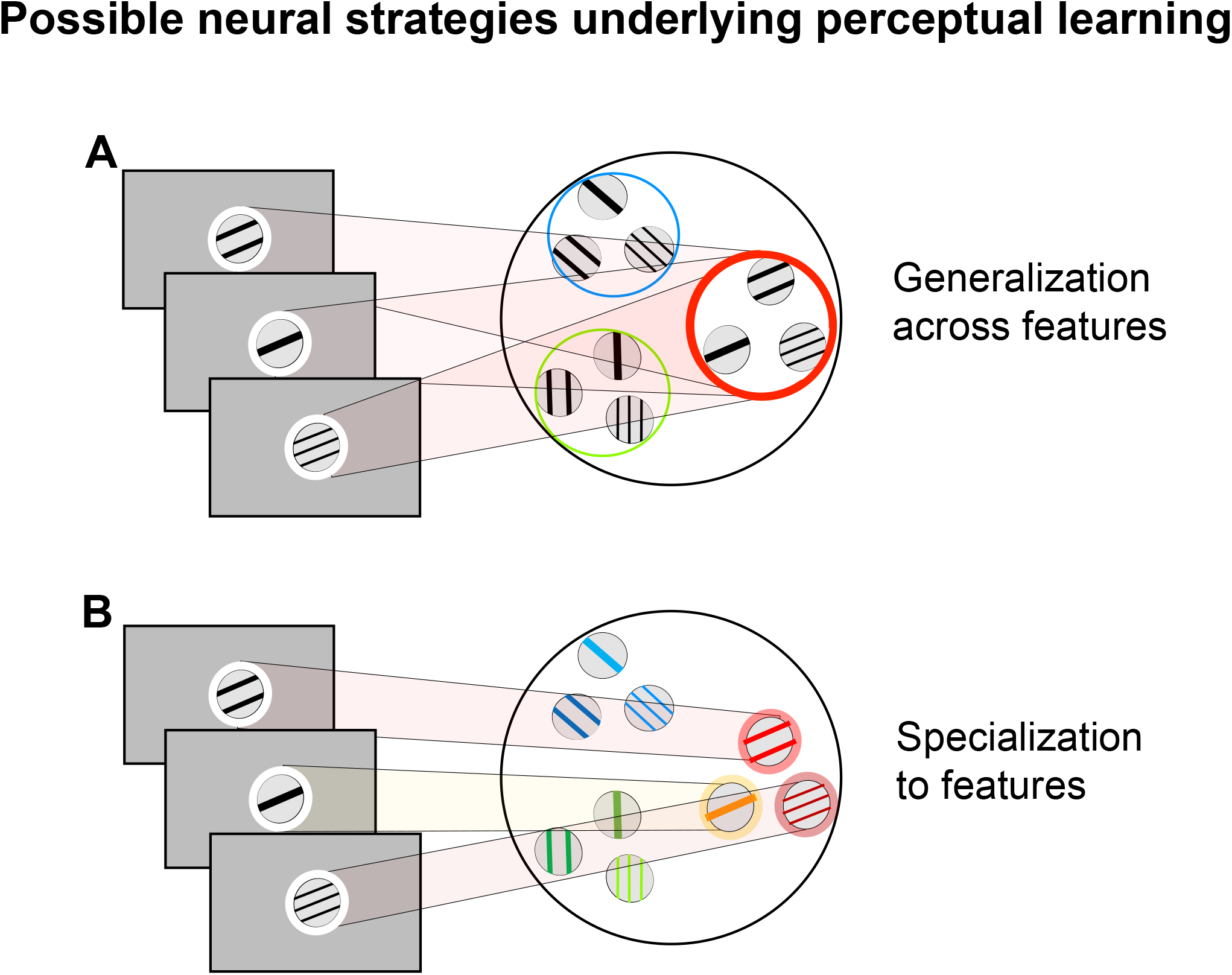
Alternative strategies for visual perceptual learning with variability. **A**. Generalization strategy: if input varies from trial to trial, learning could rely on neurons invariant to task-irrelevant features of the stimuli. These neurons deal with variability by subsuming it in their broad tuning and can accommodate generalization to new stimuli. However, their tuning to the task-relevant feature may not be very precise. **B**. Specialization strategy: alternatively, the system could implement learning with neurons narrowly tuned to task-relevant and task-irrelevant features. These neurons can provide high precision information for the task at hand. However, more neurons need to undergo plasticity than in A if inputs vary. Moreover, a specialization strategy does not lead to generalization, as new, untrained neurons are needed to accommodate new inputs.

Alternatively, the visual system could specialize on both task-relevant and -irrelevant aspects of the stimuli, using highly precise neurons narrowly tuned to both dimensions (Fig. 1B). This strategy is optimal when there is little variability because PL can be concentrated on the most informative neurons. When facing high variability, this strategy implies that separate neurons need to be recruited for each stimulus alternative during training. E.g., if oriented stimuli vary in spatial frequency (SF) during orientation discrimination PL, each orientation-SF band combination needs to be learned separately. This strategy readily assures that task performance relies on the most precise orientation information available. Yet, by relying on narrowly tuned neurons, this strategy comes at a cost, because these neurons cannot, by themselves, support generalization to other stimuli or locations in the visual field.

Here, we test how the visual system deals with the challenge of variability while achieving high performance in PL. We chose two features which are known to entail highly specific PL effects, orientation and SF^13^. During training, we systematically vary the required precision of the orientation discrimination task; in addition, subjects are trained with different degrees of variability in a task-irrelevant dimension, SF. We find that task-irrelevant SF variability indeed leads to better generalization of orientation discrimination performance to new SFs that were never shown during training, even in difficult tasks. Furthermore, subjects trained with variable stimuli can generalize better to new, untrained locations. Together, this suggests that they rely on SF-invariant neurons with large receptive fields. We then perform the same experiments in a deep neural network (DNN) that recapitulates several known PL phenomena^14^. We find a similar pattern of results, suggesting that variability-induced catalysis of generalization holds in vivo as much as in silico. We go on to show that SF variability during training leads to the recruitment (or emergence) of SF-invariant representations – and not to an increase of SF-specialized units – highlighting the benefits of invariant representations for generalization.

## Results

We trained four groups of subjects (**n**=28) in an orientation discrimination task for several days, in which they had to determine whether a grating was tilted clockwise or counterclockwise from a reference on every trial. Two groups learned a high precision version of the task, with orientation differences between 0.5 and 2.75 deg, while two other groups learned a low precision version of the task, with orientation differences between 3 and 5.25 deg. One group from each precision level was trained with a single SF (1.70 cpd), referred to as ‘low variability’ from here on, while the other group received training with three pseudo-randomly interleaved SFs (0.53, 1.70 and 2.76 cpd), referred to as ‘high variability’.

All four groups showed statistically significant increases in orientation discrimination performance as a function of training: on average, orientation thresholds decreased by 1.36° in the four training groups (Fig. 2). The initial thresholds did not differ between the high and the low variability groups in the high precision (mean difference=0.70, permutation test, *p*=0.063, Hedges’*g*=1.10) or the low precision (mean difference=0.51, permutation test, *p*=0.480, Hedges*’g*=0.46) conditions. This suggests that variability did not negatively affect task performance at the outset of training.

**Figure 2:**
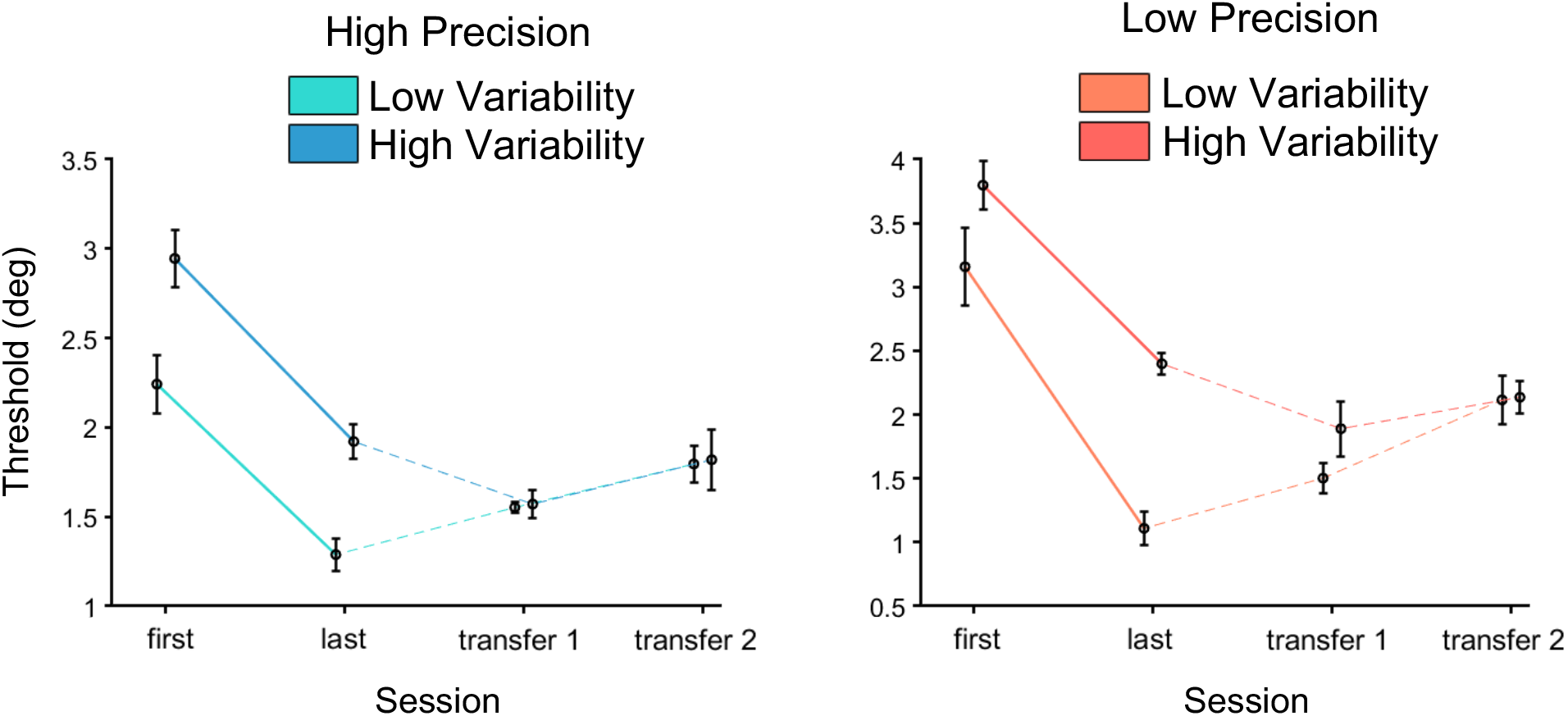
Learning curves in human subjects. To plot learning curves, we determined discrimination thresholds (using Weibull fits) per group and session. In the high precision regime, both groups show significant learning effects on thresholds (low variability: mean(last-first)=-0.95, permutation test, *p*=0.008, Hedges’ *g*=-1.68; high variability: mean (last-first)=-1.02, permutation test, *p*=0.008, Hedges’ *g*=-1.84), which do not differ between groups (mean difference between groups (last-first)=-0.07, permutation test, *p*=0.822, Hedges’ *g*=-0.13). Furthermore, the low variability training regime shows specificity of learning as its threshold increases significantly for the spatial frequency transfer (mean (transfer1-last)=0.26, permutation test, *p*=0.047, Hedges’ *g*=0.74) and again when the location is changed (mean (transfer2-last)=0.51, permutation test, *p*=0.016, Hedges’ *g*=1.49). For high variability, we find a significant reduction in thresholds with the new spatial frequency (mean (transfer1-last)=-0.35, permutation test, *p*=0.047, Hedges’ *g*=-0.72) suggesting further learning, and generalization for the location transfer (mean (transfer2-last)=0.10, permutation test, *p*=0.531, Hedges’ *g*=0.22). In the low precision regime, both groups show significant learning (low variability: mean (last-first)=-2.04, permutation test, *p*=0.008, Hedges’ *g*=-1.79; high variability: mean (last-first)=-1.4, permutation test, *p*=0.008, Hedges’ *g*=-1.85). The two groups do not differ statistically in how much they improve (mean between groups (last-first)=0.65, permutation test, *p*=0.181, Hedges’ *g*=0.75). For the low variability regime, thresholds increase numerically but not significantly with a new spatial frequency (mean (transfer1-last)=0.39, permutation test, *p*=0.063, Hedges’ *g*=0.57), but reveal specificity when the location is altered (mean (transfer2-last)=1.01, permutation test, *p*=0.016, Hedges’ *g*=1.81). The high variability group shows generalization with the new spatial frequency (mean (transfer1-last)=-0.51, permutation test, *p*=0.078, Hedges’ *g*=-0.85) and at the new location (mean (transfer2-last)=-0.26, permutation test, *p*=0.172, Hedges’ *g*=-0.37). Overall, precision, variability, and session are significant factors in explaining thresholds (precision main effect *F*(1,24)=4.384, *p*=0.047, partial *η*^2^ =0.15; variability main effect *F*(1,24)=6.732, *p*=0.016, partial *η*^2^ =0.21; session main effect *F*(3,72)=48.847, *p*<0.001, partial *η*^2^ =0.70). Furthermore, we find a significant interaction between variability and session (*F*(3,72)=7.616, *p*<0.001, partial 7 <0.20), but not between precision and session (*F*(3,72)=2.103, *p*=0.107, partial *η*^2^ =0.14). Error bars reflect the standard error of the mean corrected for within factors ^35,36^ to aid graphical interpretation of the within-subjects learning effects.

Next, we assessed how variability affects learning itself. To this end, we computed the “Learning Index” (LI)^15^, which quantifies learning relative to the baseline performance level. The average improvement in LI was 0.13 (Fig. 3A), and there were no significant differences in LIs between the groups (precision main effect *F*(1, 24)=0.32, *p*=0.574, partial *η*^**2**^=0.01; variability main effect **F**(1, 24)=0.03, *p*=0.863, partial *η*^**2**^=0.02; interaction effect *F*(1, 24)=0.05, *p*=0.830, partial *η*^**2**^=0). This suggests that variability did not negatively affect the amount of PL of the trained task.

**Figure 3:**
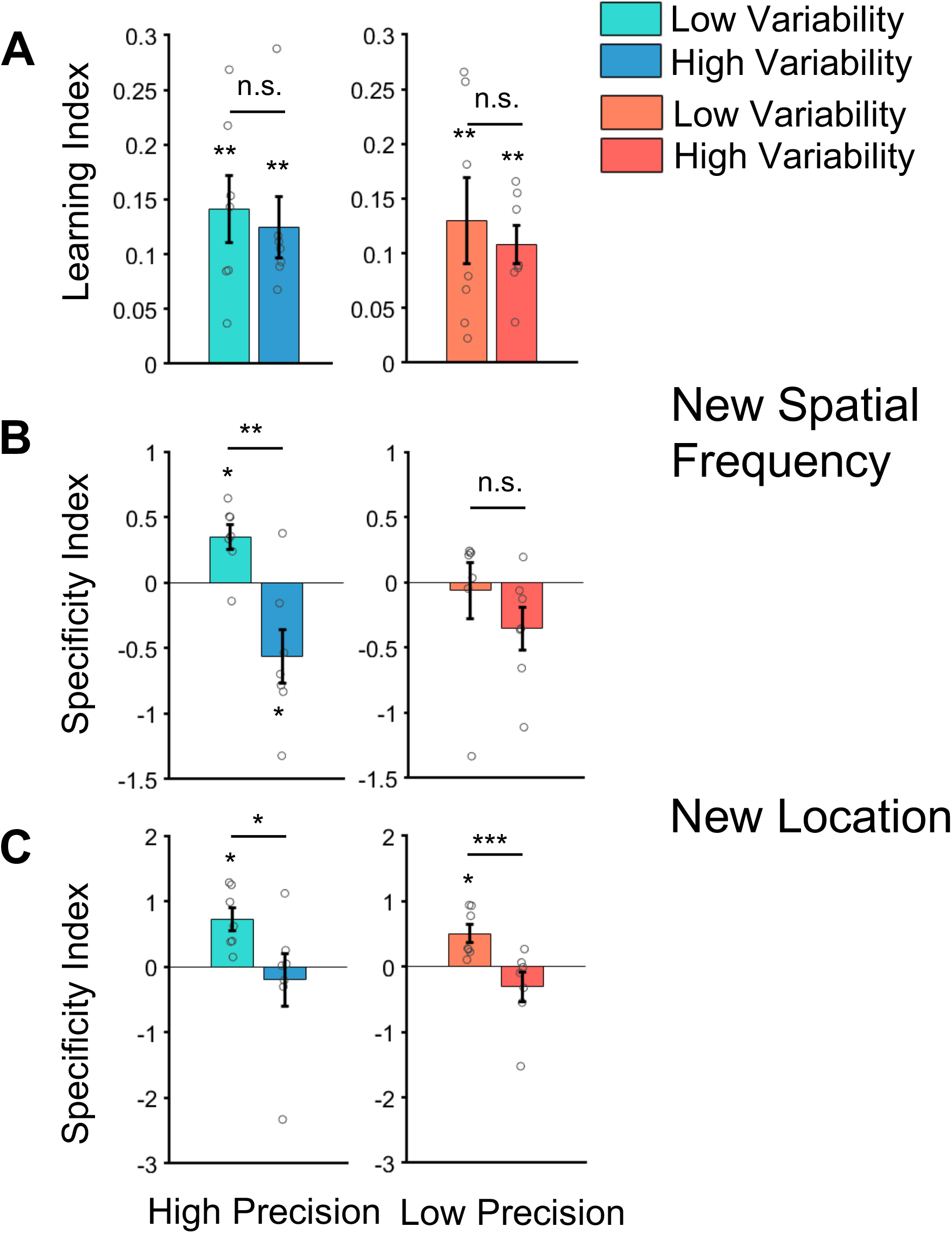
Learning and generalization in human subjects. **A.** Learning in orientation discrimination, quantified as Learning Index (LI), was significant in all groups (high precision groups; low variability mean LI=0.14, permutation test, *p*=0.008, Hedges’ *g*=2.31; high variability mean LI=0.12, permutation test, *p*=0.008, Hedges’ *g*=2.23; low precision groups; low variability mean LI=0.13, permutation test, *p*=0.008, Hedges’ *g*=1.66; high variability mean LI=0.11, permutation test, *p*=0.008, Hedges’ *g*=3.06). Task-irrelevant variability did not negatively affect the amount of learning (high precision groups mean difference in LI=0.017, exact permutation test, *p*=0.702, Hedges’ *g*=0.22; low precision groups mean difference in LI=0.022, exact permutation test, *p*=0.615, Hedges’ *g*=0.27). We obtained the same results when we quantified learning on the basis of orientation discrimination thresholds (Fig. 2) and when we trained with a different range of spatial frequencies (1.70, 2.54 and 2.76 cpd) in the high precision, high variability group (Supplemental Figs. S2 and S2). *B*. To test the specificity of learning, a new SF was presented to the subjects after the last training session. After high precision training, learning was specific for the group trained with low variability (mean SI=0.35, permutation test, *p*=0.031, Hedges’ *g*=1.81). In contrast, subjects that were trained on the same high precision task but with varying SFs fully generalized to the new SF (mean SI=-0.56, permutation test, *p*=0.047, Hedges’ *g*=-1.38). Accordingly, there was a statistically significant difference between low and high variability training within the high precision group (mean difference in SI=0.91, exact permutation test, *p*=0.004, Hedges’ *g*=2.16). Both low precision groups showed generalization (low variability mean SI=-0.06, permutation test, *p*=0.969, Hedges’ *g*=-0.15; high variability mean SI=-0.35, permutation test, *p*=0.078, Hedges’ *g*=-1.10), with no difference between them (mean difference in SI=0.29, permutation test, *p*=0.317, Hedges’ *g*=0.58). We obtained the same results when we quantified specificity on the basis of orientation discrimination thresholds (Fig. 2) and after training with a different range of spatial frequencies (1.70, 2.54 and 2.76 cpd) in the high precision, high variability group (Supplemental Figs. S1 and S2). **C**. We also tested whether subjects could perform the transfer task at a new location. Here, training variability shaped specificity and generalizability of learning similarly in the high and low precision regimes (low variability mean difference in SI=0.92, exact permutation test, *p*=0.033, Hedges’ *g*=1.14; high variability mean difference in SI=0.81, permutation test, *p*<0.0001, Hedges’ *g*=1.65): low variability groups both show specificity (mean SI 0.72 and 0.50, respectively; permutation test, *p*=0.016, Hedges’ *g*=2.12, and *p*=0.016, Hedges’ *g*=1.84). In contrast, training with high variability led to generalization (mean SI -0.20 and -0.31, respectively; permutation test, *p*=0.703, Hedges’ *g*=-0.25, and *p*=0.203, Hedges’ *g*=-0.69). We obtained the same results when we quantified specificity on the basis of orientation discrimination thresholds (Fig. 2). In all panels, error bars reflect the standard error of the mean, circles reflect individual subjects. *** stand for *p*<0.001, ** for *p*<0.01, and * for *p*<0.05.

We then tested generalization of PL in the four groups. For analyses, we computed the “Specificity Index” (SI,^16^), which quantifies how much of the learning improvement can be carried over to previously untrained conditions. Positive SI values indicate that learning is specific, i.e., does not transfer to new conditions, whereas SI values smaller than or equal to 0 indicate generalization.

We first challenged subjects with a new, untrained SF (0.96 cpd) outside the SF channels of the closest trained SFs. A 2 × 2 analysis of variance (ANOVA) of the four SIs revealed a statistically significant interaction between the factors precision and variability (Fig. 3B, *F*(1, 24)=4.52, *p*=0.044, partial *η*^*2*^=0.11): PL in the high precision, low variability group was highly specific to the trained SF (Fig. 3B left, mean SI=0.35, permutation test, *p*=0.031, Hedges’ *g*=1.81), as would be expected from a classical PL training paradigm with only a single SF. In contrast, subjects that were trained on the same high precision task but with varying SFs fully generalized to the new SF and in fact showed negative SI (mean SI=-0.56, permutation test, *p*=0.047, Hedges’ *g*=-1.38), implying that they even continued to improve their performance in the transfer task. We obtained similar results when we compared thresholds instead of SIs (Fig. 2). To rule out that generalization to 0.96 cpd was due to the bracketing of this new SF by two trained SFs (0.53 and 1.7 cpd) in the high variability group, compared to only one SF (1.7 cpd) in the low variability group, we additionally performed the same experiment in a new group of subjects trained at 1.7, 2.54, and 2.76 cpd. Here, variability also led to better generalization than low variability in the absence of bracketing (Supplemental Figs. S1B and S2). The results from high precision training thus suggest that task-irrelevant variability indeed enables generalization in PL.

In the low precision group, a different picture emerged (Fig. 3B right). Here, both low and high variability training led to good generalization with SIs not significantly different from 0 (low variability mean SI=-0.06, permutation test, *p*=0.969, Hedges’ *g*=-0.15; high variability mean SI=-0.35, permutation test, *p*=0.078, Hedges’ *g*=-1.10), and no statistically significant difference between both training regimes (mean SI difference=0.29, exact permutation test, *p*=0.317, Hedges’ *g*=0.58). We again obtained similar results when we compared thresholds instead of SIs (Fig. 2). This result is in accordance with previous PL studies showing that easy tasks generalize well^16^ (but see below).

The results from the SF transfer condition suggest that high variability and low precision both enable generalization. Given that subjects trained with high precision or low variability could perform the task in a SF band that lay outside the trained SF range is indicative of a strategy involving SF-invariant neurons. These neurons are more prevalent in higher order visual cortex, where neurons also have larger receptive fields^17^. We thus hypothesized that if subjects relied on invariant neurons (and not many narrowly tuned neurons which are more prevalent in early visual areas), they should also show transfer to new spatial locations. This is because trained and untrained locations would be covered by the same receptive fields. We thus moved the stimuli to a new, iso-eccentric location in the same quadrant 8 dva from the original training location and repeated the transfer task there.

Here, we find that all groups trained with only one SF cannot transfer their learning gains to the new location, irrespective of the required precision in the orientation discrimination task (mean SI 0.72 and 0.50, respectively; permutation test, *p*=0.016, Hedges’ *g*=2.12, and *p*=0.016, Hedges’ *g*=1.84). In contrast, both groups that were trained with variable SFs were able to generalize (mean SI -0.20 and -0.31, respectively; permutation test, *p*=0.703, Hedges’ *g*=-0.25, and **p**=0.203, Hedges’ *g*=-0.69). This was also evident in terms of a main effect of variability in the ANOVA (Fig. 3C, *F*(1,24)=23.08, *p*<0.0001, partial *η*^*2*^*=*0.32; all other *p*>0.27, partial *η*^*2*^<0.02). These results suggest that learning with variability indeed taps on SF-invariant neurons with larger receptive fields than classical PL with only a single SF band.

To gain further insight into the computations underlying generalization after high variability training, we repeated the same experiments in a DNN purpose built for PL^14^. This network, which is derived from the general AlexNet architecture^18^, recapitulates several known behavioral and physiological PL effects, e.g., higher learning rates for low precision tasks^16^ and the sharpening of tuning curves as a result of training that has originally been observed in the primary visual cortex of non-human primates^19^.

We trained this network with our four training paradigms, crossing the factors precision and variability, as in our human subjects. Each condition was simulated 25 times over 360 training steps. When considering the final performance on the trained task, we again find statistically significant learning effects in all four groups (Fig. 4A; mean LI 1, 0.85, 1, and 0.98, respectively, all *p*<0.0001, all *t*(24)>72.36, all Hedges’ *g*>14.01). Yet, in contrast to human subjects, there is a significant interaction between variability and precision in LIs (*F*(1,96)=115.17, *p*<0.0001, partial *η*^*2*^*=*0.54), with a larger difference between high and low variability training in the high precision than the low precision simulations (mean difference in LI=0.138, *p*<0.0001, Hedges’ *g*=2.99). This difference notwithstanding, critically, when we challenge the network with a new, untrained SF, we observe a differential pattern of generalization performance that resembles what we found in human observers, most clearly in the high precision training regime: networks trained with variable SFs show better transfer to a new SF than networks trained with only a single SF (Fig. 4B; mean difference in SI=0.39, *t*(48)=16.03, *p*<0.0001, Hedges’ *g*=4.46). We obtain a similar result with low precision (mean difference in SI=0.64, *t*(48)=55.29, *p*<0.0001, Hedges’ *g*=15.39). Across training regimes, variability explained most between-group variance (variability main effect *F(*1, 96)=1457.58, *p*<0.0001, partial *η*^*2*^=0.94, compared to precision main effect *F*(1, 96)=32.18, *p*<0.0001, partial *η*^*2*^=0.25, and interaction *F*(1, 96)=85.01, *p*<0.0001, partial *η*^*2*^*=*0.47). Hence, while training leads to different LIs and more specific learning effects in DNNs than in humans, the overall pattern of generalization results bears resemblance between our in vivo and in silico results.

**Figure 4:**
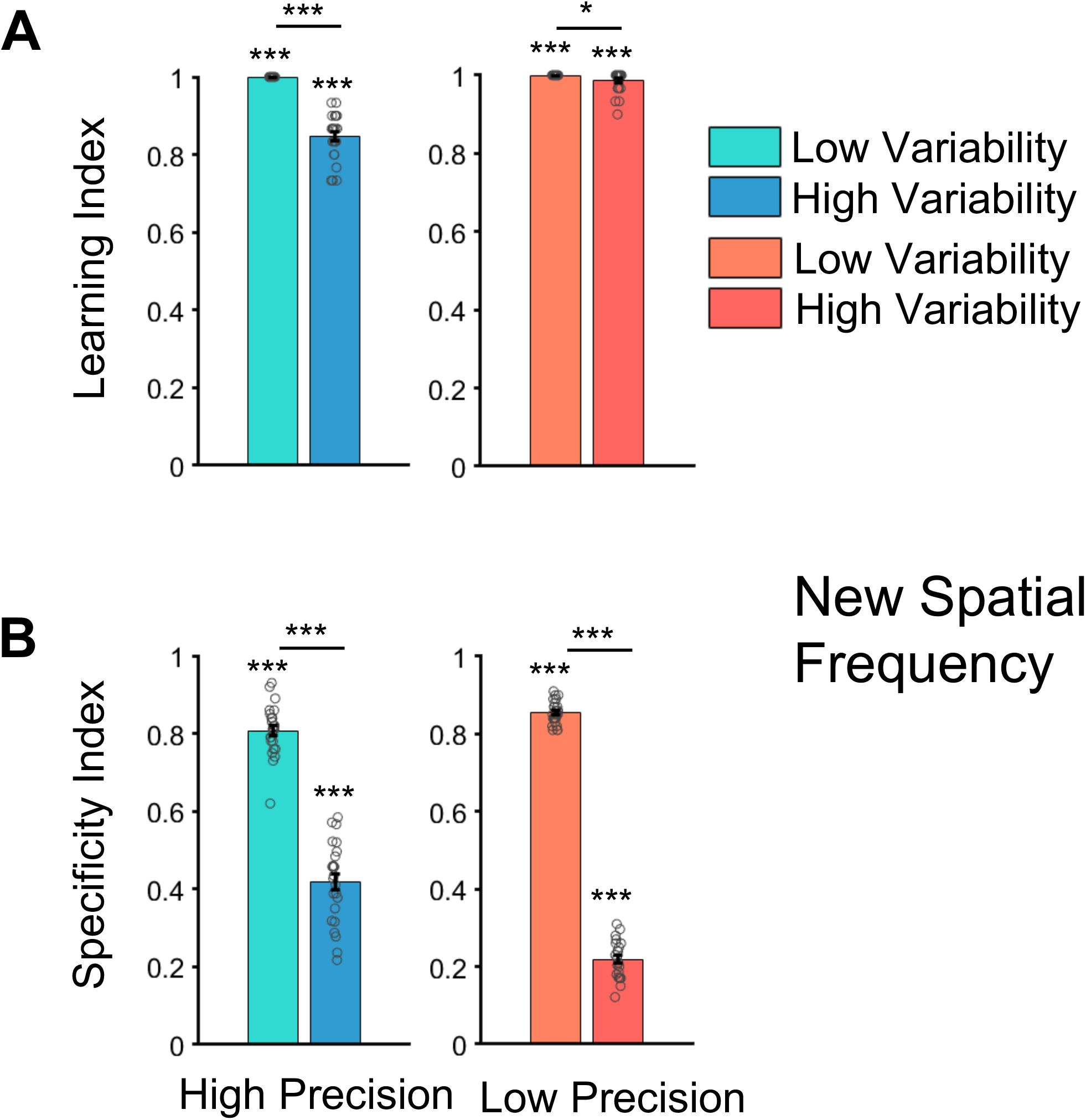
Learning and generalization in deep neural networks. **A**. We found significant learning effects across all four training regimes with high learning levels (mean LI of 1, 0.85, 1 and 0.98 respectively, all *p*<0.0001, *t*(24)>14.01). Input variability led to lower LIs than training without variability (high precision groups mean difference in LI=0.152, *t*(48)=12.97, *p*<0.0001, Hedges’ *g*=3.61; low precision groups mean difference in LI=0.013, *t*(48)=2.44, *p*=0.006, Hedges’ *g*=0.69) *B*. When the networks were fed with stimuli at a new untrained SF, we found specificity in all four groups (mean SI 0.81, 0.42, 0.85, and 0.22, respectively, all *p*<0.0001, *t*(24)>3.96). Importantly, the difference in learning specificity was strongly affected by training variability (high precision groups mean difference in SI=0.39, *t*(48)=16.03, *p*<0.0001, Hedges’ *g*=4.46; low precision groups mean difference in SI=0.64, *t*(48)=55.29, *p*<0.0001, Hedges’ *g*=15.39). In all panels, *** stand for *p*<0.001, ** for *p*<0.01, and * for *p*<0.05. Error bars reflect the standard error of the mean, circles reflect individual simulations.

To dissociate a generalization strategy relying on SF-invariant orientation representations from a specialization strategy relying on units narrowly tuned to SF and orientation, we then investigated how representations in the DNN changed as a function of training. Since the amount of training involved in the network simulations cannot easily be related to the number of trials seen by human observers, we correlated the pattern of results that we obtained with humans to the pattern of transfer results after 2, 5, 15, 35, 45, 50, 100, and 360 training steps of the networks. We find the highest correspondence between in vivo and in silico experiments after 100 training steps (correlation of SI between human subjects and DNNs, *r*=0.81, *p*<0.0001). We used the simulation results from this training step for all subsequent analyses. To compare the two strategies, we computed the number of units tuned to the trained SF and a SF-invariant orientation tuning index (SIOI) and for each of the 5 layers of the DNN (see STAR methods).

We find that instead of increasing, learning significantly decreases the number of SF-tuned across all layers (Fig. 5A and Supplemental Table S1). Hence, a specialization strategy is unlikely to explain our results. The decrease in the number of SF-tuned neurons differentiated high from low variability training in several layers (Fig. 5B and Supplemental Table S1) and was predictive of learning specificity (Fig. 5C) in layer 1 (*r*=0.85, *p*<0.0001, *R*^*2*^ =0.72), layer 4 (*r*=0.73, *p*<0.0001, *R*^*2*^ =0.53), and layer 5 (*r*=0.62, *p*<0.0001, *R*^*2*^ =0.38) in high precision training (all other layers magnitude *r*<0.02, *p*>0.210, *R*^*2*^ <0.03), and in all layers in low precision training (all *r*>0.59, *p*<0.0001, *R* >0.34). This provides a first hint at a generalization strategy. We thus turned to SIOI to further investigate the relationship between invariance and generalization. Indeed, we find that SF-invariant orientation tuning as quantified by SIOI increases as a function of training, especially in the high variability groups, with the largest effects in the top layer of the DNN (Fig. 6A, B and Supplemental Table S2). Furthermore, from layer 1 onwards, SIOI predicts generalization on the transfer task (Fig. 6C) in high precision training (layer 1 *r*=-0.30, *p*=0.035, *R*^*2*^ =0.09; layer 2 *r*=-0.88, *p*<0.0001, *R*^*2*^ =0.77; layer 3 *r*=-0.85, *p*<0.0001, *R*^*2*^ =0.72; layer 4 *r*=-0.86, *p*<0.0001, *R*^*2*^ =0.74; layer 5 *r*=-0.89, *p*<0.0001, *R*^*2*^ =0.78) and low precision training (layer 1 *r*=-0.78, *p*<0.0001, *R*^*2*^ =0.61; layer 2 *r*=-0.95, *p*<0.0001, *R*^*2*^ =0.90; layer 3 *r*=-0.98, *p*<0.0001, *R*^*2*^ =0.96; layer 4 *r*=-0.98, *p*<0.0001, *R*^*2*^ =0.96; layer 5 *r*=-0.97, *p*<0.0001, *R*^*2*^ =0.97). This suggests that variability-induced generalization in PL could indeed rely on a recruitment (or emergence) of invariant representations that provide orientation information irrespective of SF.

**Figure 5:**
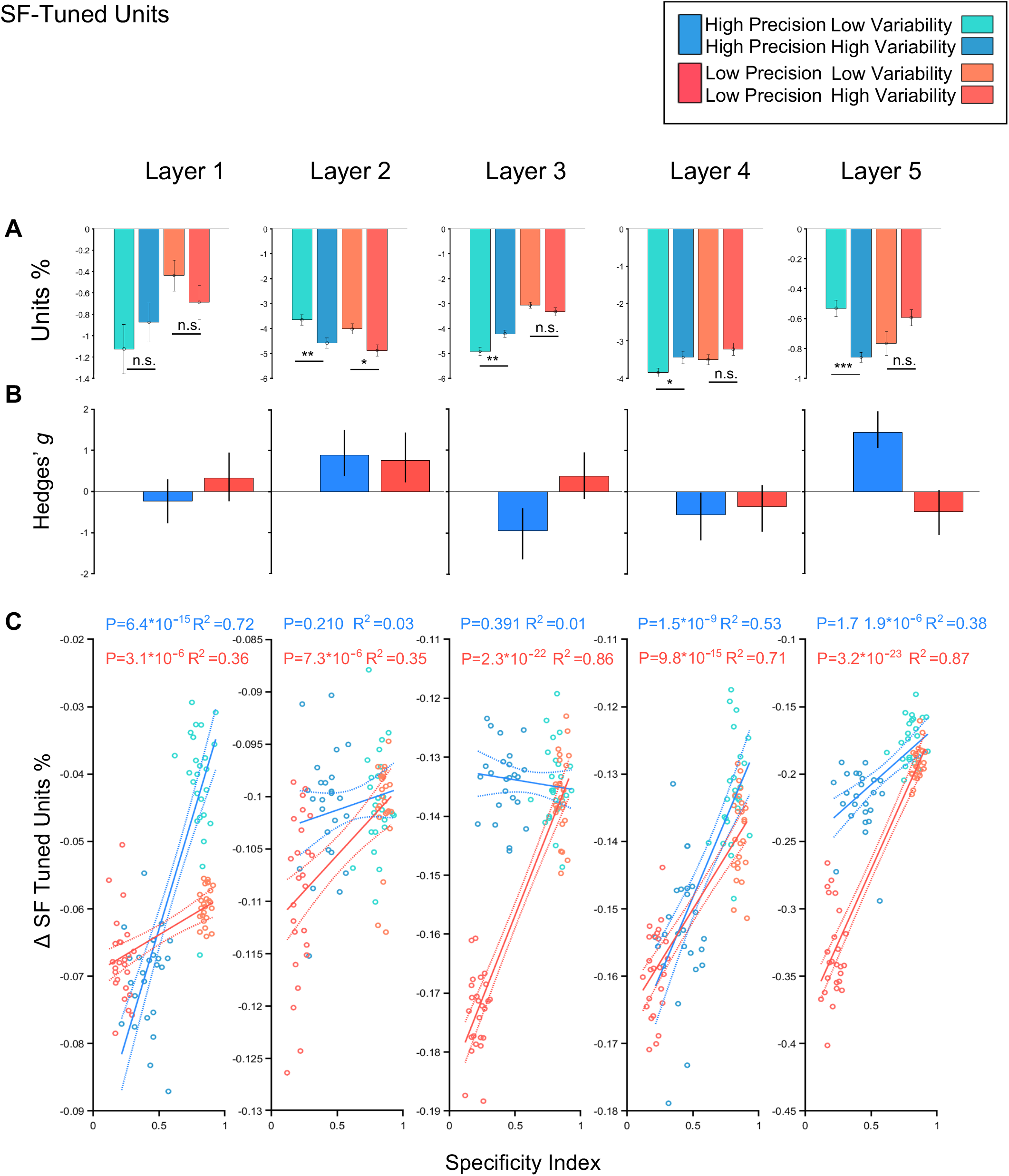
Effects of learning on the number of SF-tuned units. **A**. The number of SF-tuned units decreases significantly across all layers and all groups (all *p*<0.003). Error bars reflect the standard error of the mean. *** stand for *p*<0.001, ** for *p*<0.01, and * for *p*<0.05. *B*. The effect size for comparing the change in the number of SF-tuned units between low and high variability groups (expressed as Hedges’ *g*) varies in size and direction across layers and training regimes (0.24 and 0.32 for layer 1; 0.88 and 0.77 for layer 2; -0.94 and 0.37 for layer 3; -0.56 and -0.36 for layer 4 and 1.44 and -0.49 for layer 5). Error bars reflect the 95% confidence interval of the effect size. **C**. The change in the number of SF-tuned units is predictive of learning specificity in layer 1 (*r*=0.85, *p*<0.0001, *R*^2^ =0.72), layer 4 (*r*=0.73, *p*<0.0001, *R*^2^ =0.53), and layer 5 (*r*=0.62, *p*<0.0001, *R*^2^ =0.38) in high precision training (all other layers magnitude *r*<0.02, *p*>0.210, *R*^2^<0.03), and in all layers in low precision training (all *r*>0.59, *p*<0.0001, *R*^2^>0.34). Dotted lines reflect 95% confidence intervals.

**Figure 6:**
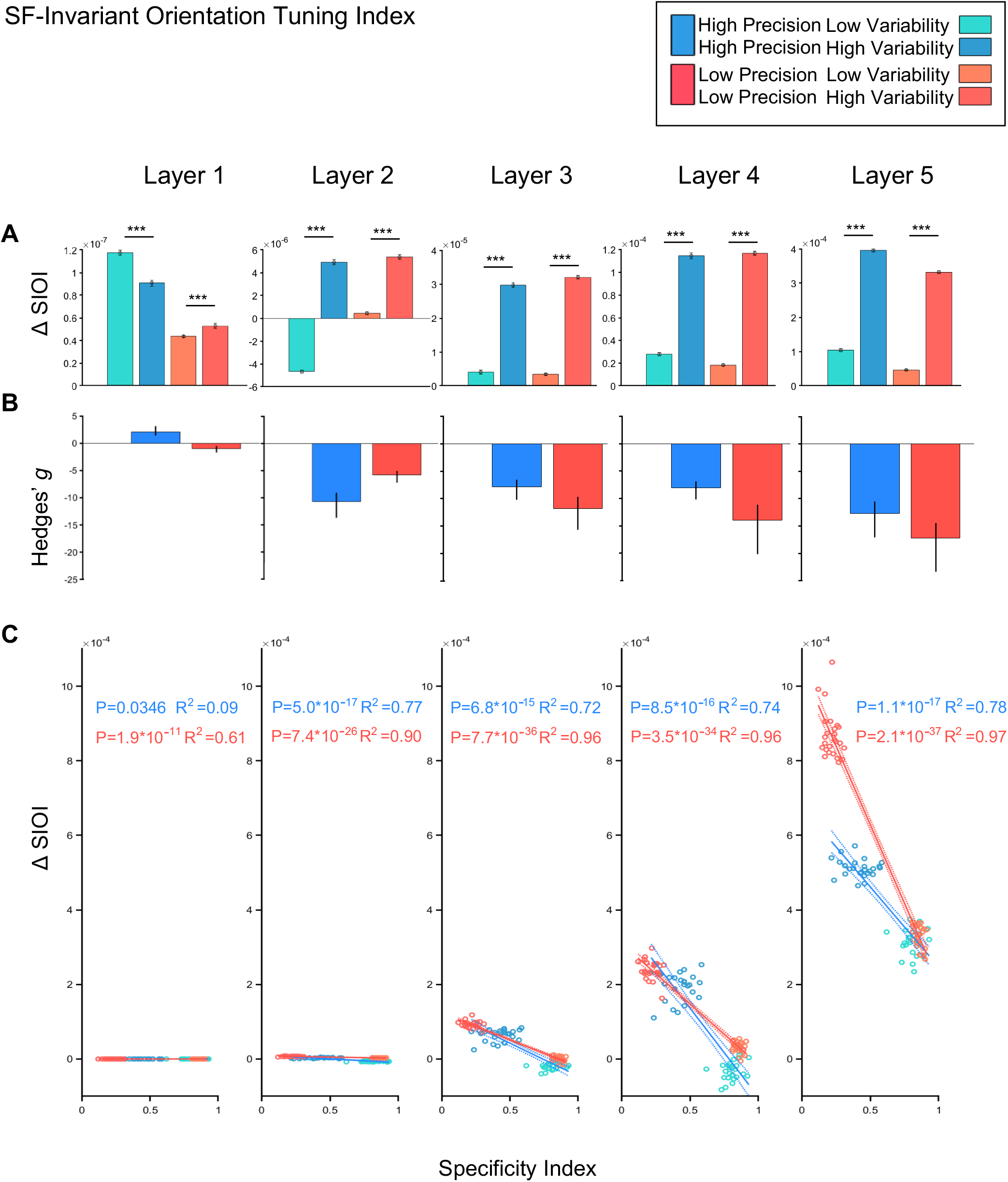
Effects of learning on SF-invariant orientation tuning. **A**. With low precision training, SIOI increases significantly (all *p*<0.0001) over training in all layers (with the exception of layer 3, where there is no change in SIOI for low variability training, *p*=0.994). This general trend is conserved for high precision, high variability training (all *p*<0.0001), while the low variability group shows significant SIOI decreases in layers 2, 3 and 4 (all *p*<0.0001). Error bars reflect the standard error of the mean. *** stand for *p*<0.001, ** for *p*<0.01, and * for *p*<0.05. **B**. The effect size for comparing the change in SIOI between low and high variability groups (expressed as Hedges’ *g*) differentiates high from low variability training in all layers for high and low precision and increases across layers (−0.48 and -2.60 for layer 1; -8.33 and -6.00 for layer 2; -6.34 and -10.91 for layer 3; -6.83 and -9.70 for layer 4 and -6.37 and -10.68 for layer 5). Error bars reflect the 95% confidence interval of the effect size. **C**. A strong negative correlation between SIOI and SI is found from layer 1 onwards (all *p*<0.035) and increases over layers. Dotted lines reflect 95% confidence intervals.

## Discussion

We find that variability enables generalization in PL, beyond the generalization benefits that have been reported in low precision tasks. Variability-induced generalization outside the trained SF band and far from the trained location overcomes two of the hallmarks of PL, namely stimulus and location specificity. Our results in human subjects align with the known tuning properties of SF-invariant neurons in the primate visual system. A role of such neurons in generalization is further suggested by our in silico results, which show that networks trained with variable stimuli evolve SF-invariant representations of orientation with training.

Traditionally, PL protocols have not involved variability, training with a minimal number of stimuli instead. This has led to many important insights, but mostly on extremely specific PL effects. E.g., simply showing stimuli to a different eye after monocular training can completely abolish learning effects^20^. By eliminating variability and presenting the same stimuli over and over, traditional PL paradigms may have unwittingly promoted ‘memorization’ or ‘overfitting’ of specific stimuli instead of generalization^11^. Variability during training endows the visual system with more robustness towards changes in the stimulus material. This also makes PL less “strange”: the high specificity of PL has not been observed in other domains of training – instead, variability has been shown to benefit transfer in domains as disparate as baseball and soccer practice^21,22^, language learning^23^, and mathematics and problem solving^24^. Hence, variability as a catalyst or enabler of generalization can be considered a principle of leaning across domains.

Our data suggest that instead of memorizing a small number of specific stimuli, the visual system uses invariant representations that provide orientation information irrespective of SF. Invariance is thought to arise from systematic pooling over feature-specific representations. When there is variability in the inputs, this entails more frequent weight updates for invariant than for feature-specific representations, because only the former participate in the task on every trial. Furthermore, relying on invariant representations suggests that fewer neurons need to undergo plasticity than in a specialization strategy, hence reducing overall metabolic cost. Invariant representations for orientation exist, e.g., in VTC, where neurons represent the orientation of simple stimuli like gratings independently of other stimulus properties such as color or spatial frequency^25,26^. Although not as frequently studied as orientation representations, e.g., in the primary visual cortex, these higher order visual neurons have been shown to be causally relevant for orientation discrimination^27^. It has also been shown electrophysiologically that orientation tuning of neurons in higher order visual cortex is less accurate than in early visual areas^12^. This is one of the bases of the Reverse Hierarchy Theory (RHT)^16^ that has previously put forward that easier tasks are learned on the basis of neurons in higher order visual cortex, whereas difficult tasks require high precision information available in early visual areas. Our results are in line with RHT, as subjects trained in the low precision task showed generalization to new SFs even if originally trained only with a single SF. However, variability during training led to overall lower SIs and extended generalization to new locations, even when the task was difficult. Hence, variability needs to be considered as an additional relevant dimension beyond task difficulty in enabling generalization of PL.

The pattern of SF and location transfer differed across the four training regimes in a way that may be informative regarding the physiological basis of the effects we observed. In particular, in the high precision groups, transfer to a new SF *and* to a new location only occurred under conditions of high variability. In the low precision group, SF transfer occurred in both the high and low variability groups, but location transfer was only evident after high variability training. This pattern of results suggests that visual PL could involve (at least) three populations of neurons depending on the training regime, namely: 1. neurons tuned to SF and location, as they are commonly found in early visual cortex; these neurons could explain the SF and location specificity in the high precision, low variability group. 2. neurons invariant to SF but not location, as they can be found at intermediate stages of visual processing; these neurons could explain concomitant transfer across SF and specificity for location in the low precision, low variability group. 3. neurons invariant to location and SF, which can be found in higher visual cortex, and which could explain transfer across SF and location in the high variability training groups across precision levels. Overall, this suggests that the contribution of lower versus higher visual areas in PL is not fixed but may depend on the variability of the training regime, which can be parsimoniously explained by a single principle, invariance (in space and SF, respectively). Future studies could test whether this also holds for other feature combinations.

Traditional PL theories have highlighted the specificity of PL^28^, but more recent studies have identified conditions other than variability under which PL can generalize (for a recent review, see ^29^). For example, in so-called ‘double training’ paradigms, practicing two tasks enables transfer across locations^30^, and in ‘training-plus-exposure’ paradigms, simultaneous or subsequent passive exposure to additional stimuli enables transfer across orientations^31^. It has also been proposed that counteracting adaptation that may arise during prolonged training can be beneficial for generalization^32^. To this end, task-irrelevant ‘dummy’ trials can be interspersed with the main task, which is akin to introducing task-irrelevant variability. We did not observe statistically significant adaptation effects in our data (Supplemental Fig. S4), possibly because we minimized stimulus-specific adaptation by randomizing phase from trial to trial. However, our experiments were not specifically designed to investigate the role of adaptation. To what extend the above-mentioned training paradigms can be understood under the same principle(s) remains an interesting and very relevant question.

Our results highlight (at least) three objectives for future studies: First, they suggest new targets for electrophysiological recordings investigating PL in non-human primates, namely higher order visual areas in VTC. Neurons in these areas code for orientation, are invariant to low-level features, and have large spatial receptive fields, and could thus support the generalization of PL we observed. Yet, studies investigating PL in these areas are, by and large, inexistent (but see^33^). Second, our results suggest that stimulus variability-induced generalization is a robust principle of learning across many learning domains, including vision, that may be strategically used in machine learning to achieve out of sample generalization. Replacing ad hoc data augmentation strategies by biologically inspired principles may lead to more robust models that may also align better with human perception. Finally, in terms of application of VPL^34^, where generalization is key, it may prove beneficial to counteract overtraining by varying stimuli in task-irrelevant dimensions.

## Acknowledgements

We would like to thank Marina Berg for help with data acquisition. This project has received funding from the European Research Council (ERC) under the European Union’s Horizon 2020 research and innovation programme (Grant agreement No. 802482, to CMS). CMS is supported by the German Research Foundation Emmy Noether Program (SCHW1683/2-1).

## CRediT author statement

G.M., Methodology, Investigation, Formal Analysis, Visualization, Data curation, Writing – original draft preparation; A.S.D., Software; C.M.S., Conceptualization, Methodology, Writing – original draft preparation, Supervision, Project administration, Funding acquisition.

## STAR METHODS

### RESOURCE AVAILABILITY

#### Lead contact

Further information and requests for resources and reagents should be directed to and will be fulfilled by the lead contact, Caspar M. Schwiedrzik.

#### Materials availability

This study did not generate new unique reagents.

#### Data and code availability

- All human behavioral data are available for download on Figshare (https://doi.org/10.6084/m9.figshare.20627052).
- The deep neural network is available from the original authors on Github (https://github.com/kevin-w-li/DNN_for_VPL).
- Any additional information required to re-analyze the data reported in this paper is available from the lead contact upon request.

### EXPERIMENTAL MODEL AND SUBJECT DETAILS

#### Human Participants

A total of 60 healthy human volunteers (34 female, 10 left-handed, mean age 28 yrs, SD 8.4 yrs) participated in this study: 37 in the main experiments (20 female, 6 left-handed, mean age 28 yrs, SD 7.2 yrs), 7 in a pilot experiment for the range of SFs (3 female, 0 left-handed, mean age 30 yrs, SD 10.5 yrs), 7 in a control experiment for potential pre-existing differences at the training and transfer location (5 female, 1 left-handed, mean age 26 yrs, SD 3.8 yrs), and 9 to test for an effect of differential bracketing of SFs (6 female, 3 left-handed, mean age 31 yrs, SD 9 yrs). All subjects had normal or corrected-to-normal vision, no neurological or psychiatric disease, and gave written informed consent before participation in accordance with the Declaration of Helsinki. Sample size was selected based on previous studies. Subjects were randomly assigned to one of four training groups (crossing the factors precision (high/low) and variability (high/low), see below). All procedures were approved by the Ethics Committee of the University Medical Center Göttingen (protocol number 29/8/17).

### METHOD DETAILS

#### General setup

All subjects were trained on a two alternative forced choice (2AFC) orientation discrimination task with oriented gratings. Training lasted 2 to 5 days with one training session per day. Transfer tasks were conducted later depending on subject availability (mean interval 2.6 days). Total training and transfer time was 4 to 7 days. Stimuli were presented on an LCD monitor (ViewPixx EEG, refresh rate 120 Hz, resolution 1920 × 1080 pixel, viewing distance 65 cm) in a darkened, sound-attenuating booth (Desone Modular Acoustics). Subjects viewed the screen through an elliptical aperture that covered the screen edges. Stimulus delivery and response collection were controlled using Psychtoolbox^37^ running in Matlab (The Mathworks, Inc.). Auditory feedback was delivered via headphones (Sennheiser HDA 280). During all experiments, we continuously acquired pupil and gaze measurements using a high-speed, video-based eye tracker (SR Research Eyelink 1000+). Data were sampled at 1000 Hz from both eyes. Subjects were paid €8 per hour. To assure constant motivation over the training sessions, subjects received a bonus of €2 if they improved by 10% from the previous training session.

#### Stimuli and task

On each trial, subjects had to decide whether a monopolar, monochromatic Gabor grating (size 3.1 dva, luminance 43.4 cd/m^2^) was tilted clockwise or counterclockwise with respect to a reference stimulus. The reference stimulus was a monopolar, monochromatic Gabor grating with identical size and constant spatial frequency (SF, 2.56 cpd) which was randomly tilted per subject, avoiding meridians and diagonals. The task stimuli were presented in ten linearly spaced difficulty levels clockwise and counterclockwise from the reference, respectively. Each condition was presented 21 times. In addition, we presented the reference orientation 21 times, amounting to a total of 441 trials per session evenly distributed among four blocks. In the high precision training groups, difficulty levels ranged from 0.5 to 2.75 deg, while in the low precision groups, orientation differences ranged from 3 to 5.25 deg. In the low variability groups, all stimuli were presented at a single SF (1.70 cpd). In the high variability groups, we used three SFs (0.53, 1.70 and 2.76 cpd). We chose these SFs such that they lie outside the other spatial frequency channels (including the one used for transfer, see below), or exactly at full width half maximum (FWHM), with the exception of 1.7 cpd which just falls into the 2.76 cpd band. For this, we assumed a SF channel bandwidth of 1.4 octaves^38^ (but see ^39^). In all groups, stimuli were presented in pseudo-random order at 12.4° eccentricity against a grey background (39.5 cd/m^2^). Phase varied randomly between 0° and 360° from trial to trial.

On each trial, we first presented the reference stimulus for 2000 ms. This was followed by a 1000 ms fixation period. Then, two grey saccade placeholders (luminance 23.54 cd/m^2^) appeared for 1500 ms, followed by the stimulus for 250 ms. After a random delay of 500-3000 ms, the choice phase started, which was indicated by a color change of the placeholders. To respond, subjects had to direct their gaze from the fixation point (size 0.31 dva, luminance 0.28 cd/m^2^) at the center of the screen to a red target (luminance 23.53 cd/m^2^) if the stimulus was rotated counterclockwise or to an isoluminant green target if it was rotated clockwise. Note that although SF and orientation are not independently processed in the visual system, we rendered SF task-irrelevant by instructing subjects to consider only orientation for their task. The assignment of colors to the placeholder locations was pseudo-randomized. Subjects had to reach the target at 12 dva distance within 1000 ms. Subjects were instructed to respond as accurately as possible. Feedback on accuracy was provided by playing a low pitch sound (incorrect) or a high pitch sound (correct) for 500 ms. The next trial started between 300 and 2000 ms later. If subjects did not respond in time or if they broke fixation (fixation window size 4.5 dva), the low pitch sound was played, and the trial was repeated later during the block. To keep subjects motivated, the high pitch sound increased in loudness after the first 2 and 3 sequential correct trials. Loudness was reset to the original level at the first incorrect trial.

Subjects were free to take breaks between blocks. There were additional breaks in the middle of each block to display feedback about performance. As a reminder, the reference was shown for 2000 ms at the center of the screen before and in the middle of each block (in total 8 times per session). Before each training session, subjects performed 16 warm-up trials.

#### Transfer conditions

Subjects had to reach at least 0.05 Learning Index (see below) improvement and show stable performance compared to the end of the preceding training session before we transitioned to the first transfer session. Subjects generally required between 2 and 5 days to reach this criterion, and there were no statistically significant differences in the amount of training days to reach criterion between the groups (*F*(3,24)=0.71, *p*=0.553, partial *η*^*2*^=0.21). For the first transfer test, we asked all subjects to perform the same task as during training, but we changed the SF of the task stimuli to a new, unseen SF (0.96 cpd). This SF was chosen to lie outside the FWHM of SF channels around the closest SFs used for training (0.53 cpd and 1.7 cpd), assuming a channel bandwidth of 1.4 cpd^38^. Pilot experiments showed that baseline performance at 0.96 cpd did not differ from baseline performance in the SFs used for training (Supplemental Fig. S4). Subjects performed 441 trials within a single transfer session. The SF of the reference remained identical. The interval between the last training session and the transfer session varied between subjects (mean 2.6 days), but there were no significant differences between the groups (*F*(3,24)=0.1, *p*=0.957, partial *η*^*2*^=0.01). Furthermore, there was no significant correlation between Specificity Indices (see below) and this interval (Pearson correlation, *r*=0.1031, *p*=0.6015). For the second transfer test, we changed the location of the transfer stimuli to a new, iso-eccentric position in the same quadrant 8 deg away from the original training location. Pilot experiments showed that there were no systematic preexisting differences in orientation discrimination performance between the training and the transfer location (mean difference in accuracy 2%, permutation test, *p*=0.28, Hedges’ *g*=0.21). Subjects again performed 441 trials within a single session.

#### Deep Neural Network Simulations

The deep learning model used in this paper was adopted from^14^. The model was implemented in PyTorch (version 1.10.0) and consists of two parallel streams, each encompassing the first five convolutional layers of AlexNet^18^ plus one fully connected layer which gives out a single scalar value. One stream accepts one fixed reference stimulus and the other stream accepts one varying target stimulus. The target stimulus is then compared to the reference stimulus. After the fully connected layers, the outputs of the two parallel streams – two scalar values - are entered to a softmax layer to give out one binary value which indicates the relative orientation (clockwise or counter-clockwise) of the target stimulus relative to the reference stimulus. We used the same feature maps and kernel size as the original paper. Each stimulus had 32×32 pixel size and was centered inside a 256×256 pixel image which was cropped to 224×224 pixel shape to make it consistent with the input size of AlexNet. All stimuli had homogenous gray background. The reference stimulus had a fixed orientation of 236° and a SF of 2.56 cpd. While keeping the same orientation and spatial frequency combination as for human subjects, we simulated four different training regimes, crossing the factors variability and precision. Each regime was independently trained 25 times. We initialized the five convolutional layers with pretrained ImageNet weights of AlexNet to mimic a (pretrained) adult brain. The last fully connected layer was initialized by zero. As for human subjects, we randomized stimulus phase for training. The training samples per group were set to batch sizes of 20 and 60 stimuli, for low variability groups and high variability groups, respectively. This ensures that each batch provides all 20 orientations and all SFs but random stimulus phases. By randomizing stimulus phase, we created 7200 and 21600 unique stimuli for the low variability groups and for high variability groups, respectively. Training parameters were set as follow: learning rate = 0.00001, momentum = 0.9, weight decay = 0.0001. The cross-entropy loss function was used as an objective function and optimized via stochastic gradient descent.

### QUANTIFICATION AND STATISTICAL ANALYSIS

All data analyses were carried out in Matlab (The Mathworks, Inc.) and R (version 4.2.1, R Core Team, https://www.R-project.org).

#### Human behavior

Human behavioral data were analyzed using (exact) permutation *t*-tests (two-sided for comparisons between groups, and one-sample tests against 0) and analyses of variance (ANOVA). Before fitting ANOVAs, data were aligned and rank-transformed^40,41^ using the ARTool package (version 0.11.1, https://github.com/mjskay/ARTool) to satisfy distributional assumptions. For *t*-tests and ANOVAs, we computed Hedges’ *g* and partial *η*^*2*^, respectively, as effect sizes. From the original sample of 37 subjects in the main experiments, 2 subjects did not complete the experiments and were thus excluded from data analysis. 3 subjects were excluded from further data acquisition after the first session because they evidently did not follow task instructions. 4 subjects were excluded during data acquisition because lack of significant learning (but there was no difference in the number of excluded subjects between high and low variability training regimes, *p*=0.6029, odds ratio = 3, Fisher exact test). The final *n* in the main experiment was thus 28 (15 female, 5 left-handed, mean age 28 yrs, SD 7.6 yrs). Accuracy was defined as the average percentage correct per session. We excluded all trials with outliers in the reaction times per subject using the estimator *Sn*^*42*^ at a threshold of 8.5. To quantify learning, we computed the Learning Index (LI)^15^, as follows:

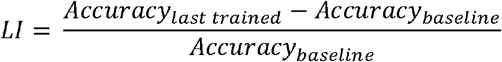

where baseline accuracy was obtained by averaging the performance of the first session. Because LI could not be lower than 0, we used one-sided (exact) permutation **t**-tests for comparisons against 0. To quantify transfer, a Specificity Index (SI)^16^ was calculated, as follows:

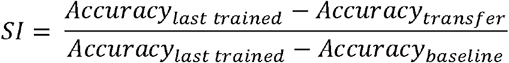

In addition, we fitted psychometric functions (Weibull) using the Palamedes Toolbox^43^ to derive orientation discrimination thresholds per subject and session.

#### Deep Neural Networks

For each simulation, learning and transfer performances were quantified by computing LI and SI as specified above. Baseline accuracy was obtained after the first training session. To investigate how the different training conditions affected stimulus representations in the network, the trained networks were evaluated with a set of stimuli covering all 20 orientations and all three SFs. We then computed dissimilarities between the average channel activities across all pairwise stimulus combinations using abs(1-Pearson correlation) for each channel pre and post training, respectively. Channels were identified as SF-sensitive if the sum of the lower triangular matrix was bigger than zero. Furthermore, we used the average channel activity to generate a representational similarity matrix (RSM) for each of the five AlexNet layers. To quantify SF-invariant orientation tuning, we computed the SF-invariant orientation tuning index (SIOI)^44,45^ by dividing the mean correlation along the off-diagonals of this RSM by the mean correlation of all other elements (but the main diagonal). To relate the pattern of generalization results in humans to that of the deep neural networks, we first computed representational dissimilarity of SIs separately for humans and deep neural networks. This step resulted in a 4×4 representational dissimilarity matrix for humans and networks, respectively. In these matrices, each dimension represents a given training regime, providing us with how dissimilar SIs were in the four training regimes. Then, we compared the representational geometries of SIs between humans and networks using Pearson correlation. All statistical analyses were carried out using parametric **t**-tests (one-sided for LI and changes in SIOI, two-sided for all others), ANOVA, and Pearson correlation.

