## Supplemental Material for "Variability in training unlocks generalization in visual perceptual learning through invariant representations"

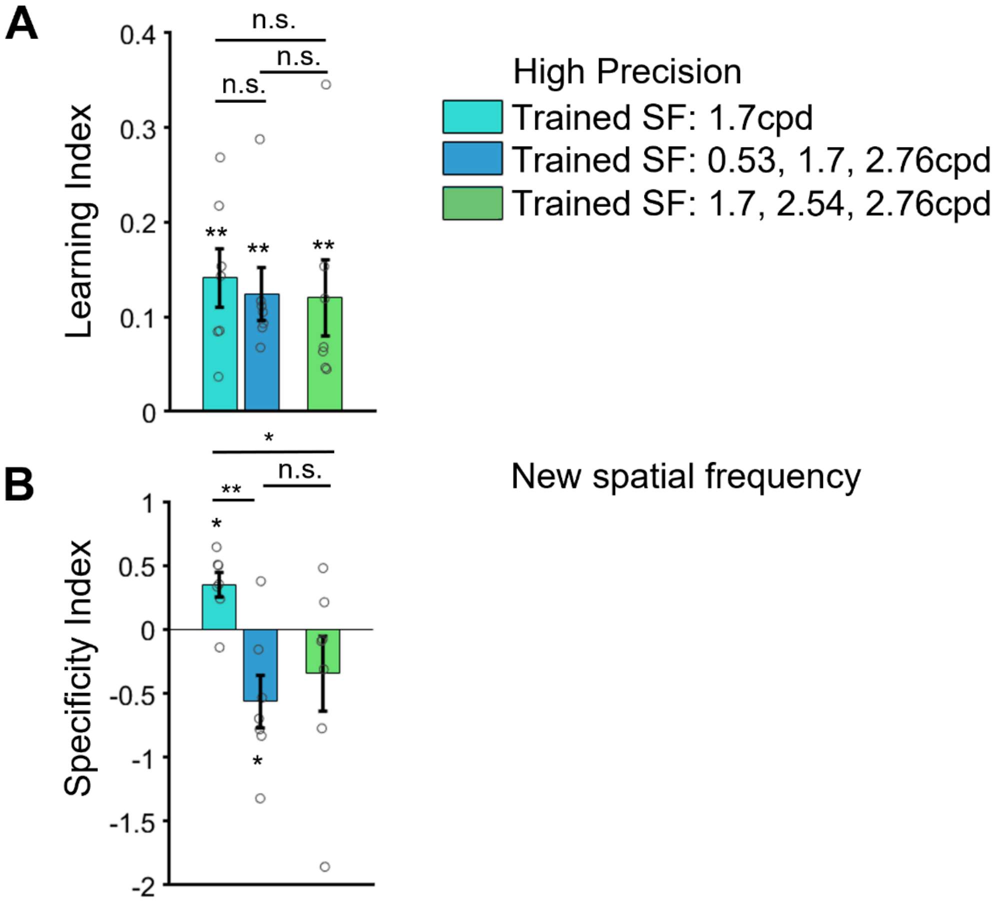


**Figure S1: Control for bracketing. Related to Figure 3.** In our main experiments, the transfer spatial frequency at 0.96 cpd was bracketed by two spatial frequencies (0.53 and 1.70 cpd) in the high variability training groups, and only bordered by one spatial frequency (1.70 cpd) in the low variability group. To rule out that differential bracketing of the transfer spatial frequency in the low versus high variability training groups could affect our transfer results, we conducted a control experiment with a different set of spatial frequencies for high precision, high variability training. Specifically, we dropped 0.53 cycles/deg as a training spatial frequency and replaced it by 2.54 cycles/deg. Hence, we trained with three different spatial frequencies (1.70, 2.54 and 2.76 cpd). All other parameters (stimulus size, eccentricity, duration, etc.) were otherwise identical to the main experiment. Nine subjects (6 female, 3 left-handed, mean age 31 yrs, SD 9 yrs) participated in this experiment. 2 Subjects were excluded because they did not complete data acquisition (final $n$=7, 5 female, 2 left-handed, mean age 33 yrs, SD 9.1 yrs). **A.** As in the main experiment, we found significant learning effects (mean LI=0.12, permutation test, $p$=0.0078, Hedges' $g$=1.48). The amount of learning did not differ significantly from the high precision, low variability group (mean difference in LI=0.021, permutation test, $p$=0.683, Hedges' $g$=0.22) nor from the original high precision, high variability group (mean difference in LI=0.004, permutation test, $p$=0.913, Hedges' $g$=0.05) **B.** When the new high precision, high variability group was tested with stimuli at an untrained spatial frequency (0.96 cpd), we found no specificity (mean SI=-0.35, permutation test, $p$=0.156, Hedges' $g$=-0.59). Importantly, the new high precision, high variability group showed more generalization than the high precision, low variability group (mean difference in SI=0.69, exact permutation test, *p*=0.026, Hedges' *g*=1.20), but did not differ significantly from the original high precision, high variability group (mean difference in SI=-0.22, permutation test, *p*=0.560, Hedges' *g*=-0.33), fully replicating the results from the main experiment. Hence, differential bracketing cannot explain the difference between high and low variability training. In all panels, *** stand for *p*<0.001, ** for *p*<0.01, and * for *p*<0.05. Error bars reflect the standard error of the mean, circles reflect individual subjects.

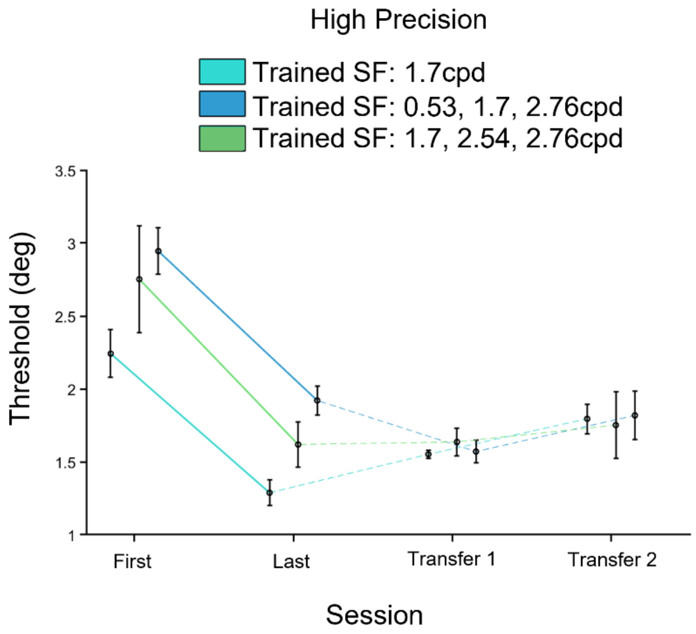


**Figure S2: Thresholds for bracketing control. Related to Figure 3.** We determined orientation discrimination thresholds (using Weibull fits) per group and session. In the high precision regime, all groups show significant learning effects on thresholds (low variability: mean(last-first)=-0.95, permutation test, *p*=0.008, Hedges' *g*=-1.68; high variability: mean (last-first)=-1.02, permutation test, *p*=0.008, Hedges' *g*=-1.84, control group: mean (last-first)=1.13, permutation test, *p*=0.008, Hedges' *g*=1.05), which do not differ between low and high variability regimes (mean difference with original group (last-first)=-0.07, permutation test, *p*=0.822, Hedges' *g*=-0.13; mean difference with control group (last-first)=-0.18, permutation test, *p*=0.826, Hedges' *g*=-0.175 ) and neither between high variability groups (mean difference with control group (last-first)=-0.11, permutation test, *p*=0.914, Hedges' *g*=-0.11). In the transfer tests, the control group shows transfer to the new SF (mean(last-first)=-0.02, permutation test, *p*=0.854, Hedges' *g*=-0.04) and location (mean(last-first)=-0.13, permutation test, *p*=0.547, Hedges' *g*=-0.38), similar to the original high variability group. Hence, differential bracketing cannot explain the difference between high and low variability training. Error bars reflect the standard error of the mean corrected for within factors[S1, 2] to aid graphical interpretation of the within-subjects learning effects.

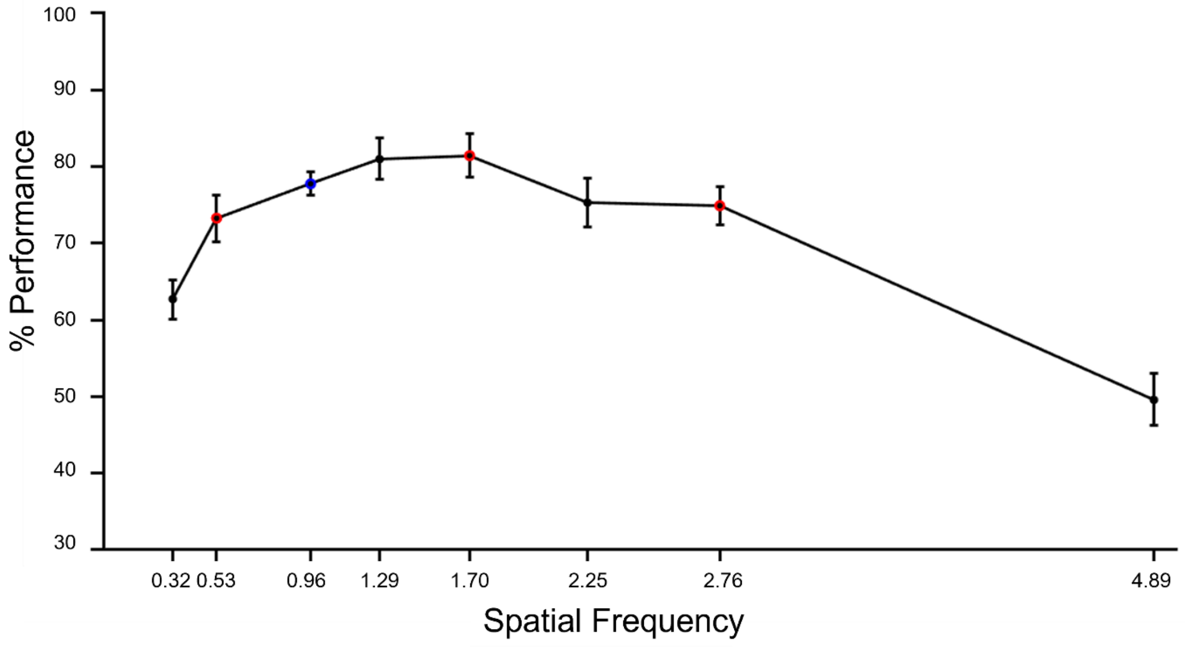


**Figure S3: Choice of transfer spatial frequency. Related to STAR methods.** We sought to find a spatial frequency to test for transfer to new spatial frequency bands that did not a priori differ significantly from the spatial frequencies used for training. To identify this spatial frequency, seven subjects (3 female, 0 left-handed, mean age 30 yrs, SD 10.5 yrs) participated in a pilot experiment. We presented oriented gratings at eight different spatial frequencies (0.32, 0.53, 0.96, 1.29, 1.70, 2.25, 2.76, and 4.89 cpd). All parameters (stimulus size, eccentricity, duration, etc.) were otherwise identical to the main experiment. We found that orientation discrimination performance at 0.96 cpd did not differ significantly from performance at the individual spatial frequencies used for training (all $p > 0.13$), nor from the mean performance across the spatial frequencies used in high variability training (mean difference 0.01, $t(6) = 0.532$, $p = 0.614$). Red data points indicate the spatial frequencies subsequently used in low (1.70 cpd) and high variability (0.53, 1.70, 2.76 cpd) training, and the blue data point the spatial frequency used in the transfer tasks (0.96 cpd). Error bars reflect the standard error of the mean.

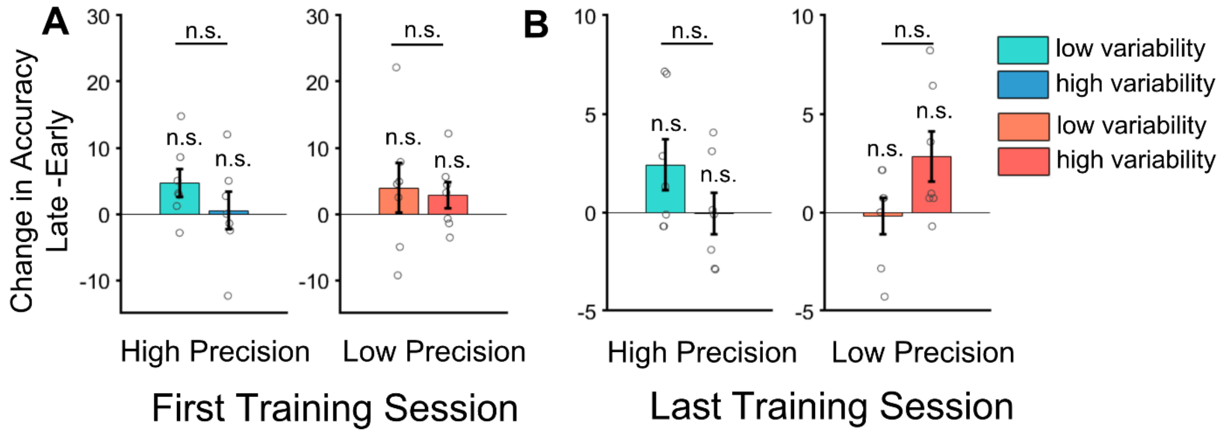


**Figure S4: Control for adaptation effects. Related to Discussion.** To assess whether there were any adaptation effects in our paradigm despite this manipulation, we quantified within-session differences in orientation discrimination performance similar as in Harris et al.[S3]. They had observed a gradual improvement in orientation discrimination performance, followed by a rebound to lower performance levels towards the end of the training session. We first quantified whether any of the groups showed a drop in accuracy towards the end of the first or the last training session, respectively. To this end, we compared accuracy in the last third to accuracy in the 2nd third of the first and last training session, respectively, per group, using one-sided permutation tests. **A.** All tests were non-significant in the first training session (all $p>0.586$, all Hedges' $g<0.63$). **B.** All tests were non-significant in the last training session (all $p>0.453$, all Hedges' $g<0.70$). The absence of a within-session performance drop in all groups indicates the absence of statistically significant adaptation effects in our data. We also compared the average accuracy in the first third of the session versus the second third of the session, and for the second third versus the last third of the session between high and low variability groups for the first and last training day, respectively. If adaptation played a role, we would expect differences between the high variability group (less adaptation) and the low variability groups (more adaptation). For the first interval, we found no difference between the high and low variability groups in how accuracy changed in first session (high precision: mean difference=-1.64, permutation test, $p=0.662$, Hedges' $g=-0.22$; low precision: mean difference=-1.45, permutation test, $p=0.770$, Hedges' $g=-0.17$) nor the last session (high precision: mean difference=-2.755, permutation test, $p=0.278$, Hedges' $g=-0.64$, low precision: mean difference=-0.68, permutation test, $p=0.708$, Hedges' $g=-0.21$). Similarly, for the second interval, we also found no difference between high and low variability groups in the first session (high precision: mean difference=4.18, permutation test, $p=0.267$, Hedges' $g=-0.63$; low precision: mean difference=1.14, permutation test, $p=0.805$, Hedges' $g=0.14$) nor the last session (high precision: mean difference=2.47, permutation test, $p=0.159$, Hedges' $g=0.79$; low precision: mean difference=-3.04, permutation test, $p=0.07$, Hedges' $g=-1.03$). These observations suggest that differential adaptation effects cannot readily explain the results we found. Error bars reflect the standard error of the mean, circles individual subjects. n.s. is not significant.

| **Change in SF-Tuned units (%)** | Group | Mean % change | *t*(24) | *p* | Hedges' *g* | Mean difference LV-HV | *t*(48)= | *p* | Hedges' *g* |
|---|---|---|---|---|---|---|---|---|---|
| Layer 1 | HP LV | -1.12 | -4.88 | <0.0001 | -1.36 | -0.25 | -0.85 | 0.399 | -0.24 |
| | HP HV | -0.88 | -4.80 | <0.0001 | -1.3 | | | | |
| | LP LV | -0.44 | -3.05 | 0.00027 | -0.85 | 0.25 | 1.17 | 0.247 | 0.32 |
| | LP HV | -0.69 | -4.34 | <0.0001 | -1.21 | | | | |
| Layer 2 | HP LV | -3.65 | -17.15 | <0.0001 | -4.77 | 0.94 | 3.18 | 0.003 | 0.88 |
| | HP HV | -4.58 | -22.47 | <0.0001 | -6.26 | | | | |
| | LP LV | -4.02 | -19.60 | <0.0001 | -5.46 | 0.84 | 2.77 | 0.008 | 0.77 |
| | LP HV | -4.88 | -21.13 | <0.0001 | -5.88 | | | | |
| Layer 3 | HP LV | -4.92 | -30.45 | <0.0001 | -8.48 | -0.71 | -3.38 | 0.001 | -0.94 |
| | HP HV | -4.21 | -31.55 | <0.0001 | -8.78 | | | | |
| | LP LV | -3.06 | -26.02 | <0.0001 | -7.24 | 0.26 | 1.34 | 0.187 | 0.37 |
| | LP HV | -3.32 | -21.44 | <0.0001 | -5.97 | | | | |
| Layer 4 | HP LV | -3.84 | -31.28 | <0.0001 | -8.71 | -0.41 | -2.00 | 0.051 | -0.56 |
| | HP HV | -3.44 | -21.34 | <0.0001 | -5.94 | | | | |
| | LP LV | -3.50 | -25.02 | <0.0001 | -6.96 | -0.28 | -1.31 | 0.197 | -0.36 |
| | LP HV | -3.22 | -19.74 | <0.0001 | -5.50 | | | | |
| Layer 5 | HP LV | -0.53 | -9.71 | <0.0001 | -2.70 | 0.33 | 5.18 | <0.0001 | 1.44 |
| | HP HV | -0.86 | -7.50 | <0.0001 | -7.50 | | | | |
| | LP LV | -0.77 | -9.61 | <0.0001 | -2.68 | -0.17 | -1.77 | 0.084 | -0.49 |
| | LP HV | -0.59 | -10.64 | <0.0001 | -2.96 | | | | |

**Table S1: The effects of learning on the number of SF-tuned units. Related to Figure 5.** The number of SF-tuned units in the deep neural network decreases significatively with learning. We report the change in the percentage of units per layer (units(after)-units(before)). To test for statistical significance against zero (columns 3:6), we used a one-sample one-sided *t*-test against 0. The difference between groups was computed as low variability – high variability. To test for statistical significance between groups trained with the same precision (columns 7:9), we used a two-sided *t*-test.

| **Change in SIOI** | Group | Mean | *t*(24)= | *p* | Hedges' *g* | Mean difference LV-HV | *t*(48) | *P* | Hedges' *g* |
|---|---|---|---|---|---|---|---|---|---|
| Layer 1 | HP LV | $14.98*10^{-8}$ | 17.79 | <0.0001 | 4.95 | $-0.000*10^{-3}$ | -1.71 | 0.093 | -0.48 |
| | HP HV | $5.54*10^{-8}$ | 23.70 | <0.0001 | 6.60 | | | | |
| | LP LV | $1.17*10^{-8}$ | 13.66 | <0.0001 | 3.80 | $-0.000*10^{-3}$ | -9.36 | <0.0001 | -2.60 |
| | LP HV | $3.03*10^{-8}$ | 16.88 | <0.0001 | 4.70 | | | | |
| Layer 2 | HP LV | $-6.54*10^{-6}$ | -30.69 | <0.0001 | -8.54 | $-0.001*10^{-3}$ | -29.93 | <0.0001 | -8.33 |
| | HP HV | $2.84*10^{-6}$ | 12.37 | <0.0001 | 3.44 | | | | |
| | LP LV | $2.95*10^{-6}$ | 25.1 | <0.0001 | 6.99 | $-0.004*10^{-3}$ | -21.58 | <0.0001 | -6.00 |
| | LP HV | $7.03*10^{-6}$ | 47.56 | <0.0001 | 13.24 | | | | |
| Layer 3 | HP LV | $-1.96*10^{-5}$ | -10.16 | <0.0001 | -2.83 | $-0.084*10^{-3}$ | -22.77 | <0.0001 | -6.34 |
| | HP HV | $6.46*10^{-5}$ | 20.48 | <0.0001 | 5.7 | | | | |
| | LP LV | $-1.11*10^{-8}$ | -0.00 | 0.994 | -0.00 | $-0.093*10^{-3}$ | -39.17 | <0.0001 | -10.91 |
| | LP HV | $9.26*10^{-5}$ | 51.10 | <0.0001 | 14.23 | | | | |
| Layer 4 | HP LV | $-3.03*10^{-5}$ | -5.78 | <0.0001 | -1.61 | $-0.22*10^{-3}$ | -24.57 | <0.0001 | -6.83 |
| | HP HV | $1.96*10^{-4}$ | 25.88 | <0.0001 | 7.20 | | | | |
| | LP LV | $3.14*10^{-5}$ | 10.86 | <0.0001 | 3.02 | $-0.2*10^{-3}$ | -34.82 | <0.0001 | -9.70 |
| | LP HV | $2.40*10^{-4}$ | 45.71 | <0.0001 | 12.73 | | | | |
| Layer 5 | HP LV | $3.15*10^{-4}$ | 44.40 | <0.0001 | 12.36 | $-0.1*10^{-3}$ | -22.87 | <0.0001 | -6.37 |
| | HP HV | $5.12*10^{-4}$ | 105.2 | <0.0001 | 29.29 | | | | |
| | LP LV | $3.27*10^{-4}$ | 53.91 | <0.0001 | 15.01 | $-0.54*10^{-3}$ | -38.37 | <0.0001 | -10.68 |
| | LP HV | $8.69*10^{-4}$ | 68.15 | <0.0001 | 18.97 | | | | |

**Table S2: The effects of learning on SIOI. Related to Figure 6.** SIOI increased significantly with learning across layers and groups. We report the change in SIOI with training (SIOI(after)-SIOI(before)). To test for statistical significance against zero (columns 3:6), we used a one-sample one-sided *t*-test against 0. The difference between groups was computed as low variability – high variability. To test for statistical significance between groups trained with the same precision (columns 7:9), we used a two-sided *t*-test.
